# A loss-of-function mutation in *Itgal* contributes to the high susceptibility of Collaborative Cross strain CC042 to *Salmonella* infections

**DOI:** 10.1101/723478

**Authors:** Jing Zhang, Megan Teh, Jamie Kim, Megan M. Eva, Romain Cayrol, Rachel Meade, Anastasia Nijnik, Xavier Montagutelli, Danielle Malo, Jean Jaubert

**Author notes:** These authors contributed equally to this work (Jing Zhang, Megan Teh). These authors jointly supervised this work (Danielle Malo, Jean Jaubert).

## Abstract

*Salmonella* are intracellular bacteria that are found in the gastrointestinal tract of mammalian, avian, and reptilian hosts. They are one of the leading causes of foodborne infections and a major threat for human populations worldwide. Mouse models have been extensively used to model distinct aspects of the human *Salmonella* infections *in vivo* and have led to the identification of several host susceptibility genes. We have investigated the susceptibility of Collaborative Cross strains to intravenous infection with *Salmonella* Typhimurium as a model of human systemic invasive infection. In this model, strain CC042 displayed extreme susceptibility with very high bacterial loads and mortality. CC042 mice showed lower spleen weight and decreased splenocyte numbers before and after infection, affecting mostly CD8^+^ T cells, B cells, and all myeloid populations. Uninfected mice also had lower thymus weight with reduced total number of thymocytes and double negative and (CD4^+^, CD8^+^) double positive thymocytes. Analysis of bone marrow resident hematopoietic progenitors showed a strong bias against lymphoid primed multipotent progenitors, which are the precursors of T, B and NK cells. An F2 cross between CC042 and C57BL/6N identified two significant QTLs on chromosome 7 (*Stsl6* and *Stsl7*) with WSB-derived susceptible alleles. A private variant in the integrin alpha L (*Itgal*) gene is carried by CC042 in the *Stsl7* QTL region. A quantitative complementation test confirmed the impact of *Itgal* loss of function in a (C57BL/6JxCC042)F1 background, but not in a C57BL/6J inbred background. These results further emphasize the utility of the Collaborative Cross to identify new host genetic variants controlling susceptibility to infections and improve our understanding of the function of the *Itgal* gene.

**Author summary:** *Salmonella* are one of the leading causes of foodborne infections and a major threat for human populations worldwide. Not all humans are equally susceptible to *Salmonella* infection. Some individuals will develop minor symptoms and recover while others develop severe illness and might die. Mouse models are used to study distinct aspects of human *Salmonella* infection *in vivo*. We used a new genetically diverse mouse population to investigate host susceptibility differences to *Salmonella* infection. We identified one mouse strain with an extreme susceptibility to infection characterized by very high bacterial loads and mortality. Mice of this strain had small thymus and spleen, two organs which are very important for producing a fully mature immune system. We showed that the strain’s immune response is impaired and that its extreme susceptibility to *Salmonella* infection is due to multiple genes defects. We identified a loss-of-function mutation in the *Itgal* gene (Integrin Subunit Alpha L) that plays a central role in the immune response to infection. This gene explains part of the susceptibility and other gene(s) involved remain to be identified. Our results emphasize how new genetically diverse animal models can lead to the identification of new host genetic variants controlling susceptibility to pathogens and improve our understanding of human infections.

## Introduction

*Salmonella enterica* is a relatively common Gram-negative bacteria that is generally transmitted via the consumption of contaminated food or water [1]. Infection with *Salmonella* can lead to a variety of pathologies with worldwide health and economic costs. Human restricted *Salmonella* strains *S*. Typhi and *S*. Paratyphi result in typhoid fever causing an estimated 190, 000 deaths per year and is typically observed in nations lacking adequate sanitation and clean drinking water programs [2, 3]. Symptoms of typhoid include fever, abdominal pain and general malaise [4]. In contrast, non-typhoidal strains such as *S*. Typhimurium lead to 93.8 million cases of gastroenteritis annually [5]. Symptoms of gastroenteritis involve diarrhea, vomiting and nausea [1]. In immunocompromised patients, non-typhoidal strains can also result in systemic and invasive infections involving bacteremia and sepsis [6].

Study of *Salmonella* in mouse models is typically conducted with *S*. Typhimurium as it is known to induce systemic infections in mice similar to the bacteremia observed in immunocompromised patients [1]. After systemic infection with *S*. Typhimurium, the bacteria are rapidly cleared from the bloodstream (within 2h), followed by localization of approximately 10% of the inoculum within macrophages and polymorphonuclear cells of visceral organs such as the spleen and liver where it can replicate efficiently. In order to resolve the resulting systemic infection, the host must activate a robust innate and adaptive immune response [1, 7].

Many factors are known to be involved in the clinical outcomes and the ability of the host to clear *Salmonella* infection in both humans and mouse models. Factors include the bacterial strain, the dosage of infection, and the host immune status, microbiome and genetic makeup [1, 6, 8, 9]. Host genetics is increasingly being recognized as a crucial element involved in host susceptibility to infection. While many genes such as toll like receptor 4 (*TLR4*), interleukin 12 (*IL12*) and signal transducer and activator of transcription 4 (*STAT4*) have been implicated in the *in vivo* response to *Salmonella*, the complete cohort of genes involved has yet to be determined [8, 10–12]. Nonetheless, establishing the genetic factors that influence disease is essential for the elucidation of host immune response pathways and will enable the identification of novel targets for therapeutic drug development.

One approach used for the detection of novel genes involved in complex traits such as *Salmonella* susceptibility utilizes a murine genetic reference population known as the Collaborative Cross (CC) [13]. While traditional models tend to use highly homogenous mouse populations, the CC has been designed to model the range of genetic variation of human population[14]. The CC is a panel of recombinant inbred mice derived from eight founder strains including five laboratory strains and three wild-derived inbred strains [15] resulting in highly variable phenotypes. The genomes of the CC strains feature relatively well dispersed recombination sites and balanced allele origins from all eight founder strains [16] allowing for the genetic dissection of complex trait [17]. Moreover, the CC serves as a platform to develop improved models of infectious disease and to map loci associated with variations in susceptibility to pathogens [18].

We previously utilized the CC to demonstrate that host genetic factors contribute to significant variation in *Salmonella* susceptibility [19]. Following challenge of 35 CC strains with *S*. Typhimurium we showed that the bacterial burdens of the spleen and liver were significantly different between strains [19]. One strain in particular known as CC042/GeniUnc (CC042) was shown to be extremely susceptible to *S*. Typhimurium infection with greater than 1000-fold higher colony forming units (CFU) in the spleen and liver as compared to the highly susceptible C57BL/6J reference strain [19]. The high susceptibility of the C57BL/6J strain is due to a missense mutation in the Solute Carrier Family 11 member 1 gene (*Slc11a1*) which has been inherited by the CC042 strain [19]. While the *Slc11a1* mutation partially accounts for the high susceptibility of CC042 mice, other host genetic variants are required to explain the extreme CC042 phenotype and have yet to be identified. The current work reports the characterization of the CC042 immunophenotype, the mapping of two loci associated with the susceptibility phenotype and the identification of a causal variant. CC042 mice were found to have a primary immunodeficiency with alterations in spleen, thymus and bone marrow hematopoietic cell populations. Quantitative trait locus mapping showed that the *Salmonella* susceptibility phenotype was controlled by at least two regions on chromosome 7 (*Stsl6* and *Stsl7*). Within the *Stsl7* locus, a *de novo* 15 bp deletion mutation in the intron 1 splice acceptor site of the integrin alpha L chain gene (*Itgal*) resulting in complete protein abrogation was shown to increase susceptibility to *Salmonella* infection. This study provides a foundation for investigating the role of *Itgal* in the response to *Salmonella* and illustrates how the CC can serve to identify mechanisms of infectious and immunological traits.

## Results

### Clinico-pathological characterization of CC042

To characterize the observed *Salmonella* susceptibility in CC042 mice, clinico-pathological parameters were evaluated in naïve CC042 mice where C57BL/6J mice were used as reference as they have a well-characterized immune phenotype prior and during *Salmonella* infection. While C57BL/6J mice are typically considered susceptible to infection, they were previously shown to be more resistant to *Salmonella* Typhimurium infection relative to CC042 mice [19].

CC042 did not present any other overt visible phenotypes prior to infection with the exception of a characteristic white head spot, most likely inherited from the WSB/EiJ CC founder parent [24]. Age and sex matched CC042 mice had comparable body weights to C57BL/6J mice (**Fig. 1a**). However, CC042 mice displayed significantly smaller spleens and thymi, in terms of mass, in comparison to the C57BL/6J (**Fig. 1b and 1c**). Examination of haematological parameters in naïve CC042 mice indicated lower total white blood cell and lymphocyte counts compared to C57BL/6J (**Table 1**). CC042 mice presented a small but significant increase in the number of red blood cells, hemoglobin and mean corpuscular hemoglobin concentration (MCHC) (**Table 1**). In addition, the splenic microarchitecture of CC042 mice was assessed using H&E staining and was shown to present moderate extramedullary hematopoiesis with prominent presence of megakaryocytes in comparison to C57BL/6J (**Fig. 2a**). Histopathological examination of CC042 liver, and kidney was normal (**data not shown**). The extramedullary erythropoiesis may explain the increased numbers of circulating RBCs and may be indicative of bone marrow failure.

**Figure 1.**
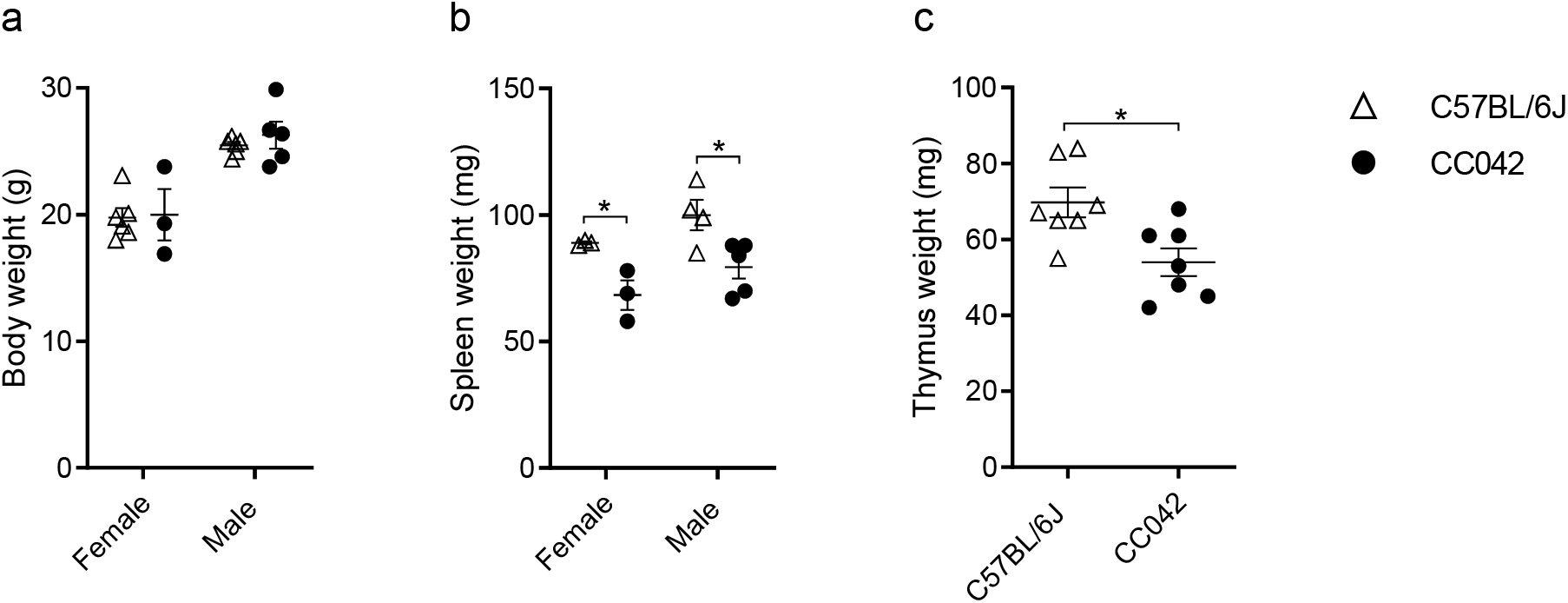
CC042 mice display reduced spleen and thymus size relative to body weight. Body (**a**), spleen (**b**) and thymus (**c**) weights for C57BL/6J and CC042 naïve mice. Only male mice were used for calculation of thymus weight and data was pooled from two experiments. Graphs represent mean ± SEM. Sidak’s multiple comparisons test (2-way ANOVA) was used to analyze body (**a**) and spleen (**b**) weights while Welch’s *t* test was used for thymus weight (**c**) where \**p* < 0.05.

**Table 1.**
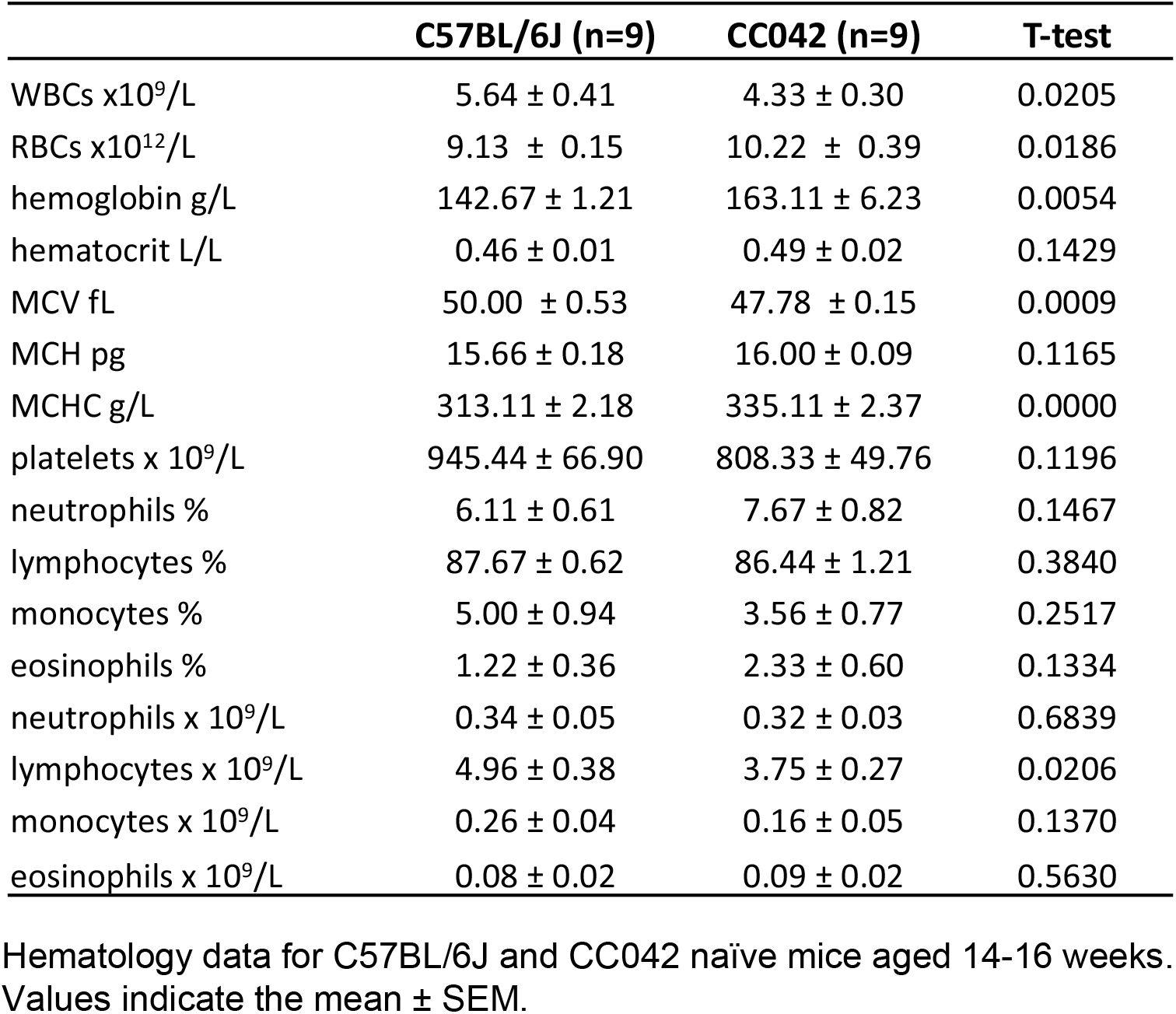
Hematologic parameters in C57BL/6J and CC042 mice

### CC042 mice display a primary immunodeficiency

To identify if the reduced splenic size (**Fig. 3a**) observed in CC042 mice may be due to alterations in the cellular immune compartment, flow cytometry for lymphoid and myeloid cells was performed on the splenocytes of CC042 and C57BL/6J naïve mice (gating scheme shown in **Supplementary Fig. S1a**). The mean total splenocyte counts in CC042 mice was significantly reduced to 31.2 ± 5.1 x 10^6^ cells compared to 75.1 ± 3.0 x 10^6^ for the C57BL/6J spleens (**Fig. 3b**). CC042 mice showed significant alterations in the splenic lymphoid compartment. Despite carrying similar numbers of splenic CD4^+^ T cells compared to C57BL/6J mice, CC042 mice displayed a reduction in the number of activated CD4^+^ T cells, as measured by CD69 expression (**Fig. 3c and 3d**). In addition, numbers of splenic CD8^+^ T cells and activated CD69 expressing CD8^+^ T cells were also significantly reduced in CC042 mice compared to C57BL/6J (**Fig. 3e and 3f**). Further CD4^+^ and CD8^+^ T cells effector function was assessed by measuring intracellular staining of IFNγ and TNFα after activation with anti-CD3 and anti-CD28. We observed diminished IFNγ and TNFα production from CC042 T cells both in naïve mice and after infection (**Fig. 3g–3j**). Reduced numbers of neutrophils, monocytes, macrophages and B cells were also observed in CC042 prior to infection (**Fig. 3k–3n**). Overall, CC042 mice present an abnormal immunophenotype characterized by reduced spleen size and generalized reduction of spleen cells affecting lymphoid and myeloid compartments. This reduction in splenic immune cells together with the observed leukopenia indicates a potential defect in leukocyte development.

**Figure 2.**
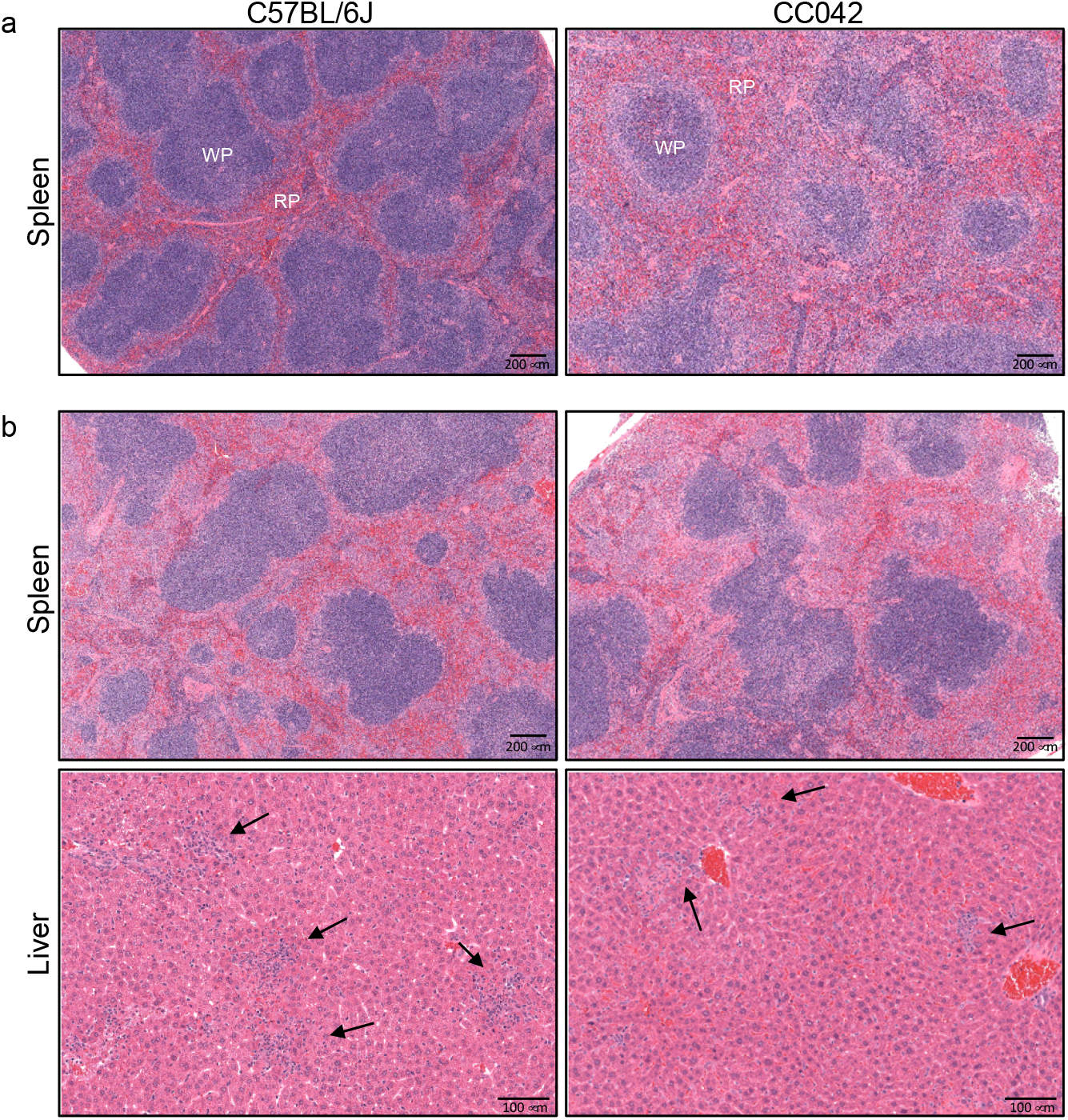
Pathologic changes in the spleen and liver of infected C57BL/6J and CC042 mice. Hematoxylin and eosin staining of C57BL/6J and CC042 spleen sections at day 0 (**a**) and spleens and liver sections at day 3 post *Salmonella* infection (**b**) representative of 6 C57BL/6J and 6 CC042 mice. Foci of necrotic hepatocytes associated with histiocytes and neutrophils were seen in both C57BL/6J and CC042 mice but were smaller and less numerous in the CC042 samples (2-4 foci per field (40X) compared to 4-5 foci in C57BL/6J). Arrows point to inflammatory foci. WP: white pulp. RD: red pulp.

**Figure 3.**
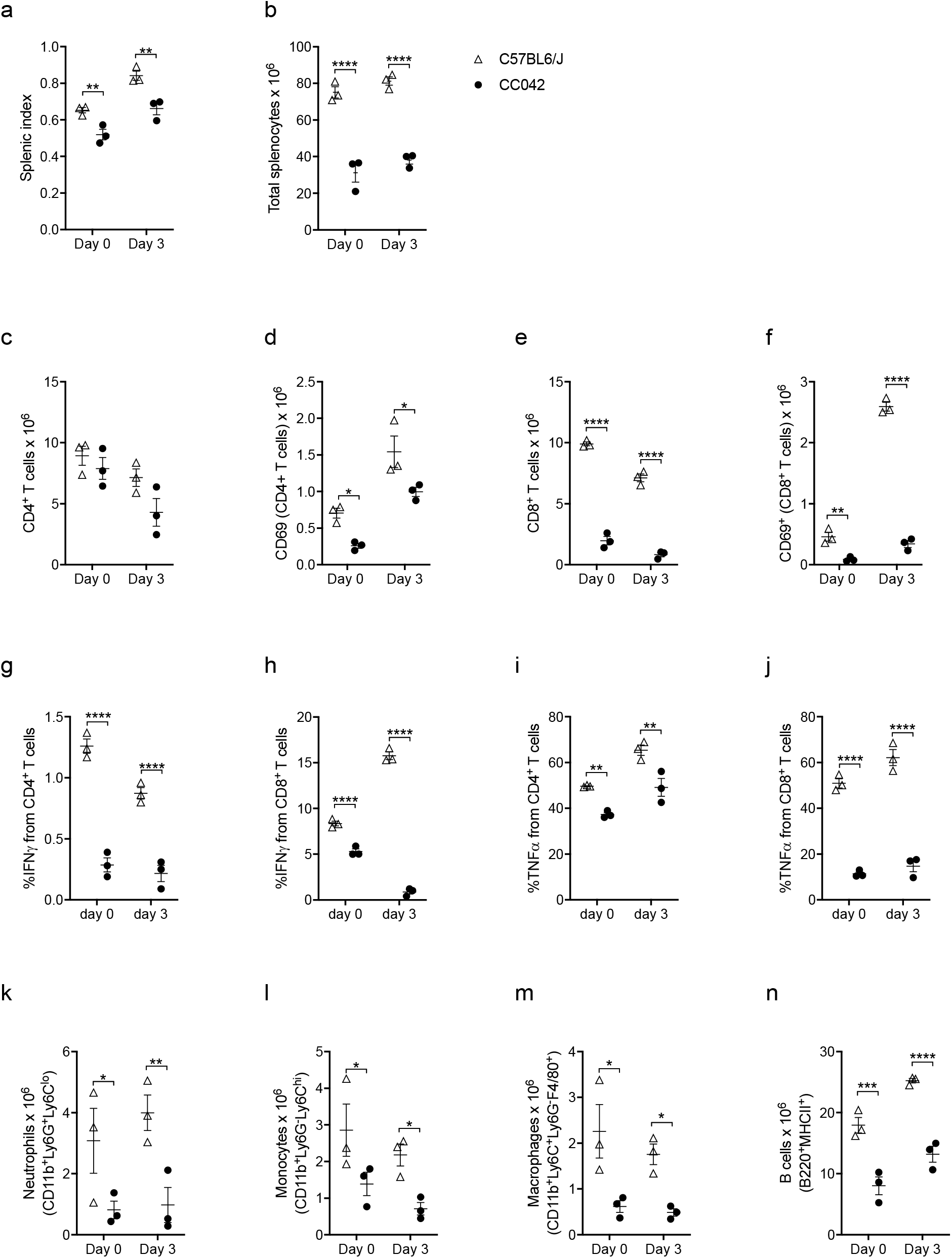
CC042 mice have significantly reduced total splenocyte numbers. Flow cytometry analysis of C57BL/6J and CC042 spleens at day 0 (naïve) and day 3 post *Salmonella* Typhimurium infection. Splenic index (**a**) and total splenocyte count (**b**) for C57BL/6J and CC042 at day 0 and day 3 of infection. Total cell counts and total CD69^+^ cells for CD4^+^ T cells (**c-d**) and CD8^+^ T cells (**e-f**). Percentage of CD4^+^ T cells and CD8^+^ T cells producing IFNγ (**g-h**) and TNFα (**i-j**) in uninfected and day 3 *Salmonella* infected splenocytes. Total cell counts for neutrophils (**k**), monocytes (**l**), macrophages (**m**) and B cells (**n**). Graphs show mean ± SEM. Data are representative of six experiments in naïve mice and three experiments at day 3 of infection. Cell populations were defined as follows: CD4^+^ T cells (TCRb^+^CD4^+^), CD8^+^ T cells (TCRb^+^CD8α^+^), neutrophils (CD11b^+^Ly6G^+^Ly6C^lo^), monocytes (CD11b^+^Ly6G^-^Ly6C^hi^), macrophages (CD11b^+^Ly6G^-^Ly6C^+^F4/80^+^), and B cells (B220^+^MHCII^+^). Analysis was conducted using Benjamini, Krieger and Yekutieli correction for multiple testing (Two-Way ANOVA) where significance is indicated as follows, \**p* < 0.05, \*\**p* < 0.01, \*\*\**p* < 0.001, and \*\*\*\**p* < 0.0001.

### CC042 mice have reduced thymic cellularity and altered T cell development

To assess if the reduction in CD8^+^ T cells and total activated T cells in the spleen of CC042 mice may potentially be due to defective T cell maturation, flow cytometry of the thymus was carried out (gating scheme shown in **Supplementary Fig. S1b**). In agreement with the previous observation that CC042 mice have reduced thymus size, CC042 mice displayed a two-fold reduction in total thymocyte counts (**Fig. 4a**). While the percentages of DP, SP CD4^+^ and CD8^+^ T cells were comparable between CC042 and C57BL/6J mice due to the reduction in thymocyte numbers in CC042 mice, a significant decrease in DN, DP, SP CD4^+^ and CD8^+^ T cells cell counts was observed in CC042 thymi (**Fig. 4b and 4c**). Further examination of the DN subset revealed significant alterations in CC042 mice (**Fig. 4d and 4e**). While no significant difference in the proportion of DN1 cells was observed, a significant reduction in the proportion of DN2 stage thymocytes was present in CC042 thymi compared to C57BL/6J controls. Unexpectedly, the diminished proportion of DN2 cells did not result in suppression of downstream subsets as a comparable fraction of DN3 thymocytes and a significantly increased proportion of DN4 cells were observed in CC042 mice compared to C57BL/6J controls. Given the reduction in total thymocytes, a significant decrease in DN1, DN2, DN3 and DN4 total cell counts was observed in CC042 mice compared to C57BL/6J controls. Overall, CC042 mice displayed a reduction in thymocyte numbers and altered progression of T cell precursors through maturation, specifically in the DN stages.

**Figure 4.**
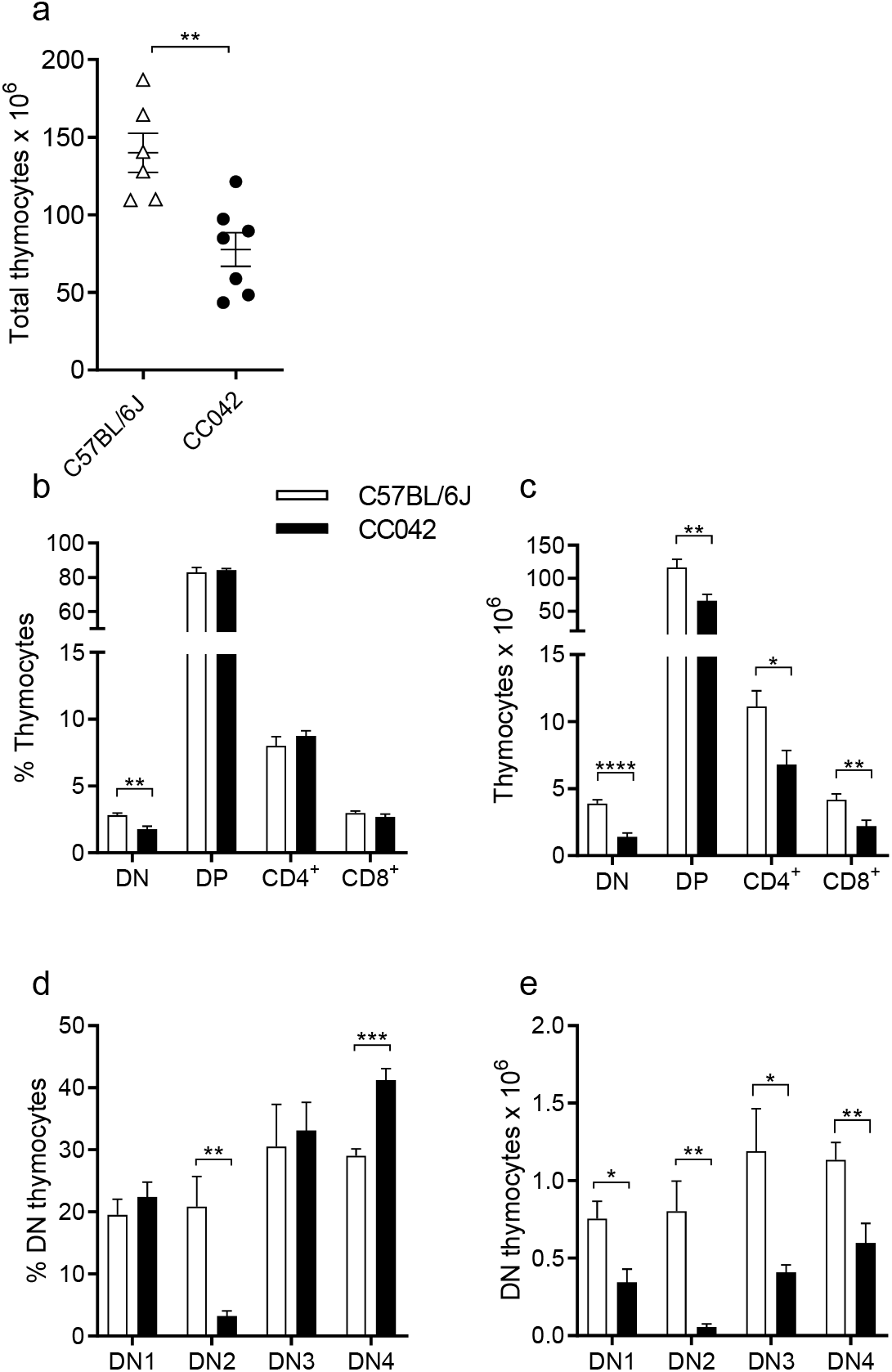
CC042 mice show significantly decreased total thymocyte numbers. Total thymocyte counts (**a**) for C57BL/6J and CC042 mice. Percentage and total thymocytes (**b-e**) by developmental stage analyzed via flow cytometry where n = 6-7 per genotype. Graphs indicate mean ± SEM with data pooled from two independent experiments. Thymocytes develop through double negative (DN) (1 to 4) to double positive (DP) stages before splitting to either CD4^+^ (CD4^+^SP) or CD8^+^ (CD8^+^SP) T cell populations. Cell populations were gated as follows: DN (CD4^-^CD8a^-^), DP (CD4^+^CD8a^+^), CD4^+^ (CD4^+^CD8a^-^), and CD8^+^ (CD4^-^CD8a^+^). DN1, DN2,DN3 and DN4 subpopulations were gated from the DN population as follows: DN1 (CD44^+^CD25^-^), DN2 (CD44^+^CD25^+^), DN3 (CD44^-^CD25^+^) and DN4 (CD44^-^CD25^-^). Multiple *t* tests using the Holm-Sidak method was used to assess significance, \**p* < 0.05, \*\**p* < 0.01, \*\*\**p* < 0.001, and \*\*\*\**p* < 0.0001.

### CC042 showed alterations in the hematopoietic progenitor populations

The observed reduction in peripheral blood leukocytes and cellularity of the spleen and thymus in CC042 mice suggested that the hematopoietic development of these populations may be impaired. To investigate haematopoiesis in CC042 mice, flow cytometry (**Table 2**) of bone marrow haematopoietic progenitor populations was conducted.

Bone marrow haematopoietic stem cells (HSCs) differentiate to multipotent progenitors (MPPs) sub-populations defined based on the expression of the CD150, CD34, CD48 and Flt3 cell surface markers (**Table 2 and Fig. 5a**) [25]. CC042 bone marrow contained less cells (**Fig. 5b**) and a reduced number of LSK (Lin^-^Sca-1^+^c-Kit^+^) cells compared to C57BL/6J with equivalent numbers of haematopoietic stem cells (HSCs) (**Fig. 5c and 5d**). However, examination of LSK subsets revealed significant alterations in the proportions of multipotent progenitors (MPP) sub-populations. The CC042 LSK compartment displayed significant reduction in downstream MPP3 and MPP4 cells in comparison to C57BL/6J controls (**Fig. 5c and 5d**). The depletion of MPP3 and MPP4 populations in CC042 bone marrow suggests a maturation block in the progression from MPP1 to MPP3 and MPP4 cell stages.

**Figure 5.**
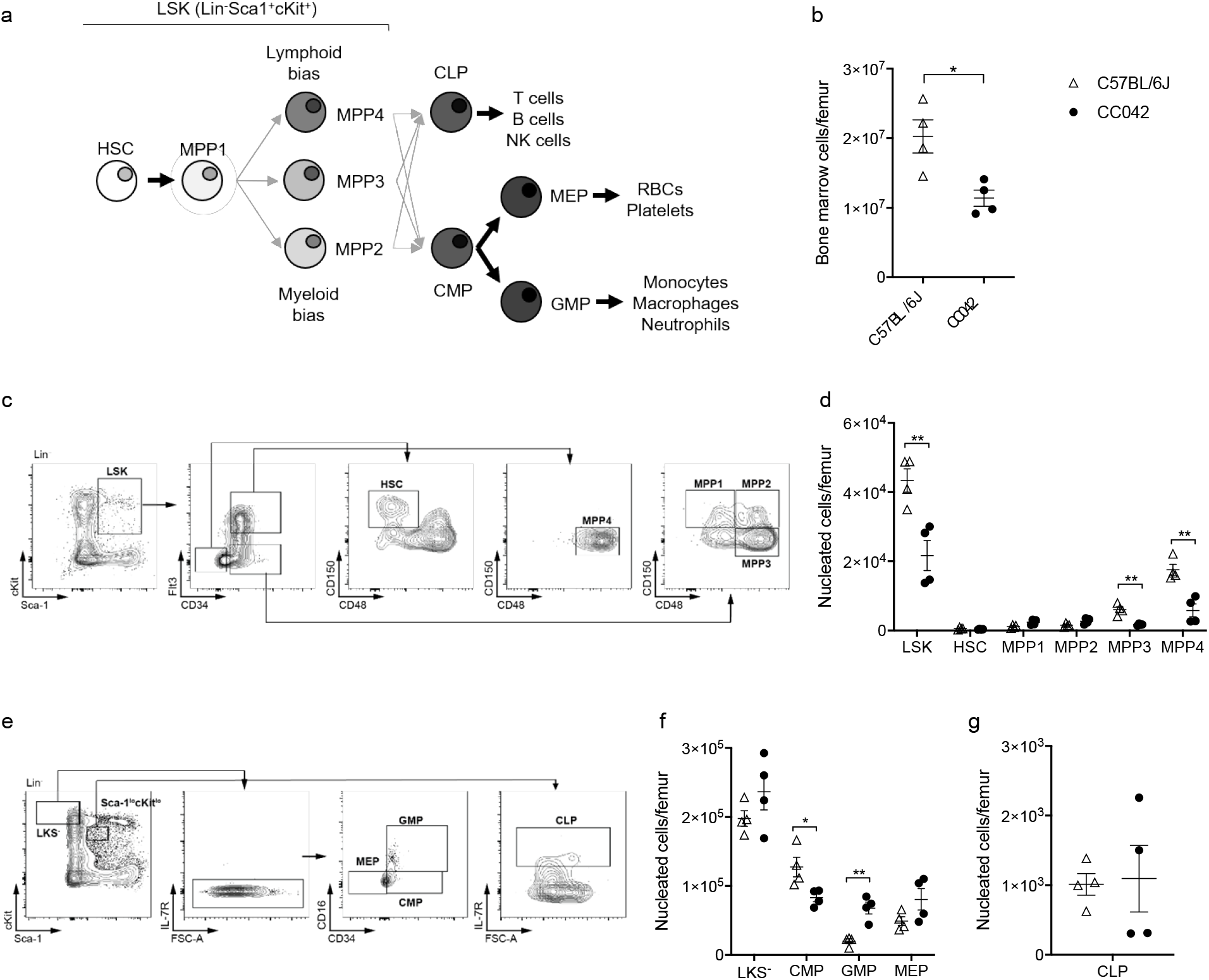
CC042 mice have altered bone marrow resident hematopoietic progenitor populations. Flow cytometry analysis of hematopoietic progenitors in femoral bone marrow from C57BL/6J and CC042 mice. Schematic diagram (**a**) of the stages of hematopoietic stem cell differentiation. Total bone marrow cells per femur (**b**) averaged over the total cells collected for two femurs. Gating scheme used to analyze Lin^-^Sca1^+^cKit^+^ (LSK) cells (**c**). Total LSK cells per femur by developmental stage (**d**). LSK cells progress through HSC, MPP1, MPP2, MPP3 and MPP4 subsets. Gating scheme used to analyze Lin^-^cKit^+^Sca1^-^ (LKS^-^) and common lymphoid progenitors (CLP) (**e**). Total LKS^-^ cells per femur grouped by progenitor stage (**f**). LKS^-^ cells comprise of common myeloid progenitors (CMP), granulocyte-macrophage progenitors (GMP) and megakaryocyte-erythroid progenitors (MEP). Total CLP per femur (**g**). Graphs show mean ± SEM. Data are representative of three independent experiments. Cell populations are defined in Table 2. Significance was determined with Welch’s *t* test for (**b**) and (**g**) and multiple *t* tests using the Holm-Sidak method for (**d**) and (**f**), \**p* < 0.05, and \*\**p* < 0.01.

**Table 2.** Hematopoietic progenitor population surface markers

| Population | Markers |
| --- | --- |
| LSK | Lin<sup>-</sup>Sca1<sup>+</sup>cKit<sup>+</sup> |
| HSC | Lin<sup>-</sup>Sca1<sup>+</sup>cKit<sup>+</sup>CD150<sup>+</sup>CD34<sup>-</sup>CD48<sup>-</sup>Flt3<sup>-</sup> |
| MPP1 | Lin<sup>-</sup>Sca1<sup>+</sup>cKit<sup>+</sup>CD150<sup>+</sup>CD34<sup>+</sup>CD48<sup>-</sup>Flt3<sup>-</sup> |
| MPP2 | Lin<sup>-</sup>Sca1<sup>+</sup>cKit<sup>+</sup>CD150<sup>+</sup>CD34<sup>+</sup>CD48<sup>+</sup>Flt3<sup>-</sup> |
| MPP3 | Lin<sup>-</sup>Sca1<sup>+</sup>cKit<sup>+</sup>CD150<sup>-</sup>CD34<sup>+</sup>CD48<sup>+</sup>Flt3<sup>-</sup> |
| MPP4 | Lin<sup>-</sup>Sca1<sup>+</sup>cKit<sup>+</sup>CD150<sup>-</sup>CD34<sup>+</sup>CD48<sup>+</sup>Flt3<sup>+</sup> |
| LKS<sup>-</sup> | Lin<sup>-</sup>cKit<sup>+</sup>Sca1<sup>-</sup> |
| CMP | Lin<sup>-</sup>cKit<sup>+</sup>Sca1<sup>-</sup>IL7R<sup>-</sup>CD34<sup>+</sup>CD16/CD32<sup>-</sup> |
| GMP | Lin<sup>-</sup>cKit<sup>+</sup>Sca1<sup>-</sup>IL7R<sup>-</sup>CD34<sup>+</sup>CD16/CD32<sup>+</sup> |
| MEP | Lin<sup>-</sup>cKit<sup>+</sup>Sca1<sup>-</sup>IL7R<sup>-</sup>CD34<sup>-</sup>CD16/CD32<sup>-</sup> |
| CLP | Lin<sup>-</sup>cKit<sup>lo</sup>Sca1<sup>lo</sup>IL7R<sup>+</sup> |

MPP sub-populations give rise to common lymphoid progenitors (CLPs) and common myeloid progenitors (CMPs) (**Fig. 5a and 5e**) [26]. CMPs form both megakaryocyte-erythrocyte progenitors (MEPs) which produce RBCs and platelets and granulocyte-macrophage progenitors (GMPs) [26]. CC042 mice showed a decreased number of CMP progenitor cells but an increase in downstream GMPs compared to C57BL/6J mice (**Fig. 5f**). No significant difference was observed in the number of CLPs between CC042 and C57BL/6J mice (**Fig. 5g**). The maturational blocks observed upstream of MPP3, MPP4 and CMP progenitors could partially explain the reduced hematopoietic cell populations present in peripheral organs.

### CC042 mice mount a distinct response to infection

To characterize the impact of infection on spleen and liver, histopathological examination of CC042 spleens and livers was conducted three days after *Salmonella* infection. The livers of C57BL/6J harbored typical lesions for this strain characterized by multifocal necrotic lesions with aggregation of histiocytes and neutrophils typical of granulomas (**Fig. 2b**). The necrotic hepatocyte foci were found to be smaller and less abundant in CC042 mice (2-4 foci per 40X field of view) than in C57BL/6J mice (4-5 foci per field of view) (**Fig. 2b**). CC042 spleens were also observed to have reduced neutrophil infiltration during infection compared to C57BL/6J spleens (**Fig. 2b**). The increased megakaryocyte numbers and erythropoiesis observed in the spleen of uninfected CC042 mice (**Fig. 2a**) was also present in the spleen of CC042 infected mice (**Fig. 2b**). During infection, CC042 mice displayed a severe splenic immunodeficiency with an important reduction in the number of activated CD4^+^ T cells, CD8^+^ T cells, activated CD8^+^ T cells, neutrophils, monocytes, macrophages and B cells compared to C57BL/6J (**Fig. 3c–3j**). The reduction in CD8^+^ T cell counts combined with the generalized defect in total T cell activation and myeloid cell numbers may account for the observed increase in *Salmonella* susceptibility.

### A mutation within Itgal is responsible for susceptibility of CC042 mice to Salmonella infection

To identify the genes which, in addition to *Slc11a1*, control the extreme susceptibility to *Salmonella* Typhimurium of CC042 mice, we produced an F2 cross between CC042 and C57BL/6NCrl, a strain closely related with C57BL/6J, with the same *Slc11a1* deficiency. The genome wide polymorphisms between the two C57BL/6 strains allowed QTL mapping even in regions where CC042 genome is of C57BL/6J origin. Infected F1 mice showed intermediate bacterial loads in the liver and spleen compared with the parental strains (**Fig. 6a and 6b**), while bacterial loads in F2 mice spanned over the parental range (see raw data in **Supplementary Table S1**). Ninety-four F2 mice with the highest or lowest liver bacterial loads were selected for QTL mapping (see raw data in **Supplementary Table S2**). Since bacterial loads in liver and spleen were strongly correlated in the 94 selected individuals (**Fig. 6c**), QTL mapping was performed on liver bacterial loads.

**Figure 6.**
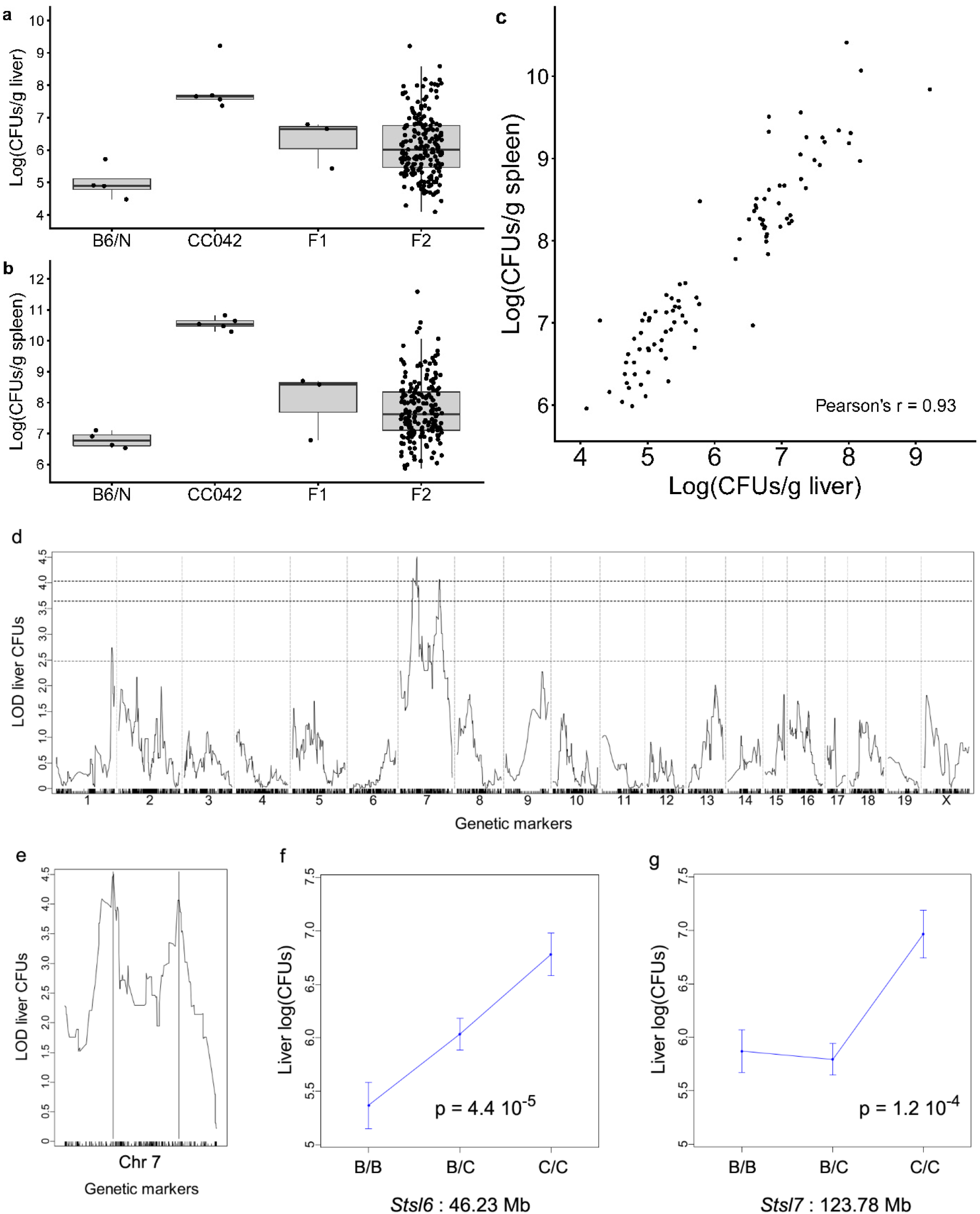
Susceptibility of CC042 mice to *S*. Typhimurium is controlled by two linked loci on Chromosome 7. Bacterial load in liver (**a**) and in spleen (**b**) at day 4 post-infection with *S*. Typhimurium in C57BL/6NCrl (n=4), CC042 (n=5), (C57BL/6NCrl x CC042) F1 (n=3) and (C57BL/6NCrl x CC042) F2 (n=196) mice. Bacterial loads in F2 mice spanned over the values of the two parental strains. Bacterial loads in the 94 individuals selected for genotyping (**c**) showing strong correlation between the two organs (Pearson’s r = 0.93). Genome-wide QTL mapping on liver bacterial load (**d**) identified two statistically significant peaks on Chromosome 7. Horizontal dashed lines indicate the 0.05, 0.1 and 0.63 (top to bottom) significance thresholds estimated from 10,000 permutations. QTL positions are indicated by vertical lines (**e**). See Table 3 for details on each QTL. The proximal *Stsl6* QTL (**f**) acted semi-dominantly on liver bacterial load, while the CC042-inherited allele at the distal *Stsl7* QTL (**g**) had a recessive mode of action. For both QTLs, the CC042-inherited allele was associated with increased bacterial load. A corresponds to the B6 allele while B corresponds to the CC042 allele.

QTL mapping identified only two significant QTLs (at 0.05 genome wide significance level), both on Chromosome 7 (**Fig. 6d and 6e**), which were named *Salmonella Typhimurium susceptibility locus-6* (*Stsl6*) and *Stsl7*. Details for each QTL are given in **Table 3**. *Stsl6* (peak position at 46.23Mb) showed semi-dominant mode of inheritance, with heterozygotes having an intermediate bacterial load compared to the two types of homozygotes (**Fig. 6f**). The CC042 allele at *Stsl7* (peak at 123.78Mb) was recessive and homozygotes had 10 times higher liver bacterial loads (**Fig. 6g**).

**Table 3.**
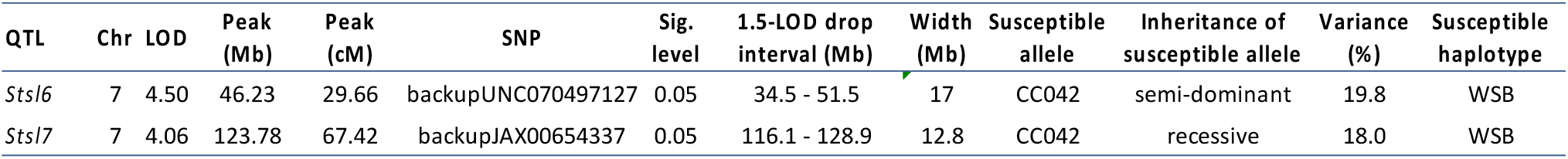
Summary of the significant QTLs identified in the (CC042 x B6/N)F2 cross

CC042 inherited *Stsl6* and *Stsl7* QTL regions from the WSB founder (**see Supplementary Fig. S2**). Other CC strains also inherited WSB alleles in either or both QTL regions but had much lower bacterial loads than CC042 (**Supplementary Fig. S2**). In particular, CC035 also carried WSB-derived alleles at *Stsl6* and *Stsl7*. This result suggested that the susceptibility alleles present in CC042 at these two loci could result from *de novo* mutations which occurred during the development of the CC042 strain. These private variants have been previously identified by the sequencing of CC strains and are publicly available [23]. Three private variants were identified in the *Stsl6* confidence interval in genes *Ush1c*, *Ccdc123*, *4930435C17Rik* and only one within *Stsl7* interval in the integrin alpha L (*Itgal*) gene, also known as *Cd11a*. This variant is a 15 base pair deletion located at the 3’ end of intron 1 and resulting in the disruption of the intron 1 splice acceptor site (**Fig. 7a**). We hypothesized that the intron 1 splice donor would attack the intron 2 splice acceptor resulting in skipping of exon 2 during splicing. Joining exon 1 to exon 3 would create a frame shift and a downstream premature stop codon, resulting in a truncated form of ITGAL protein.

**Figure 7.**
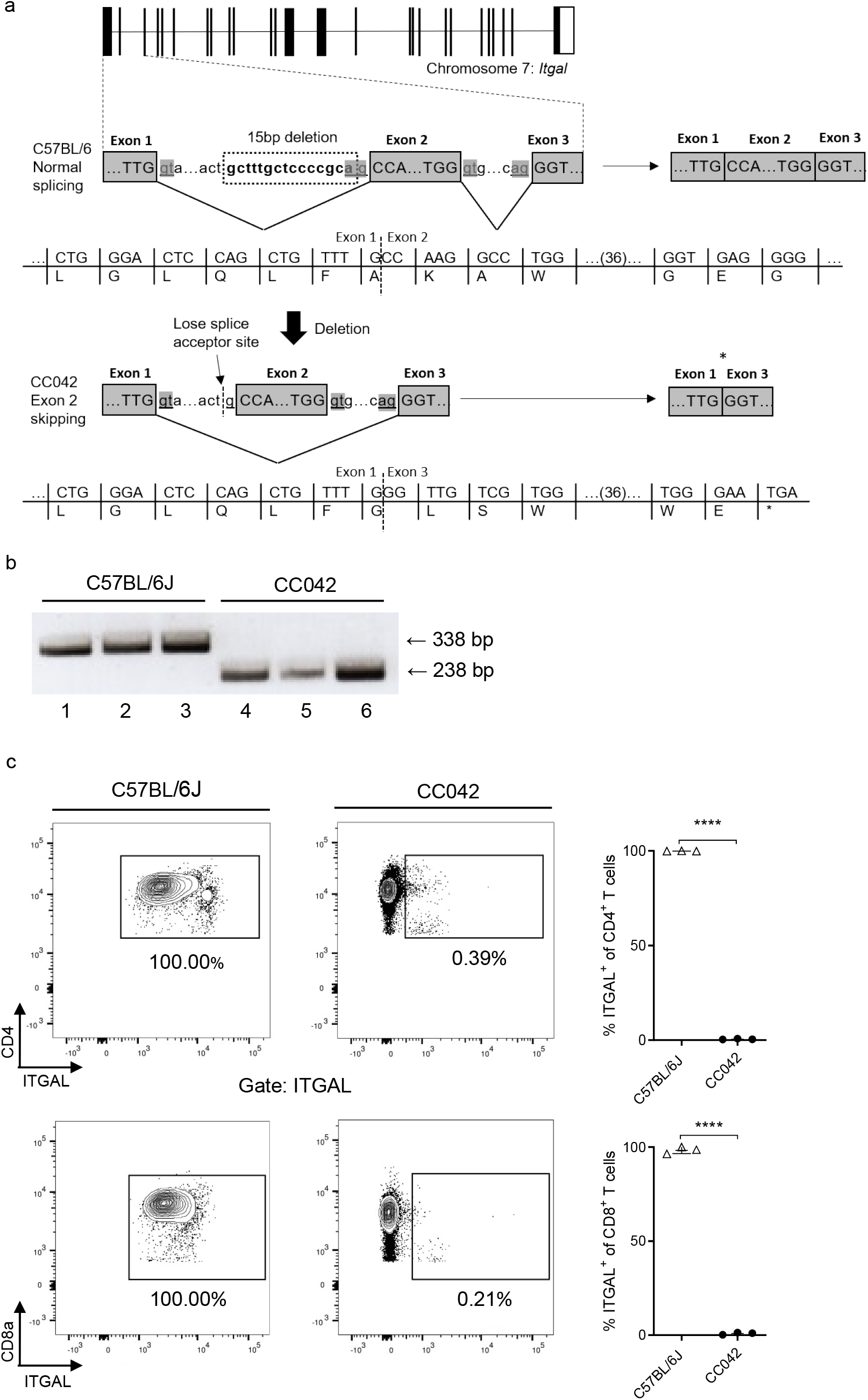
*Itgal* mutation in CC042 mice resulted in exon 2 skipping and absence of protein expression. Schematic diagram (**a**) illustrating *Itgal* mutation in CC042 mice. A 15 bp deletion results in the loss of the intron 1 splice acceptor site resulting in exon 2 skipping and the formation of a premature stop codon. PCR analysis (**b**) of *Itgal* cDNA from C57BL/6J and CC042 mice. Amplification of the region flanking exon 2 produces a 338 bp PCR product in C57BL/6J mice carrying the wild type *Itgal* gene (lanes 1-3). A 238 bp PCR product was produced in CC042 mice corresponding to a 100 bp deletion due to exon 2 skipping (lanes 4-6). (**c**) Flow cytometry analysis of ITGAL surface expression on CD4^+^ and CD8^+^ T cells. ITGAL gates were constructed using fluorescence minus one panels and significance was calculated using Welch’s *t* test, \*\*\*\**p* < 0.0001. Graphs represent the mean ± SEM. Significance was determined with Welch’s *t* test, \*\*\*\**p* < 0.0001.

To verify this prediction, reverse transcription and amplification of *Itgal* transcripts was conducted in C57BL/6J and CC042 cells using primers flanking exon 2. C57BL/6J cells were found to express a 338 bp transcript indicative of the inclusion of exon 2 (**Fig. 7b**). In contrast, CC042 cells produced a 238 bp transcript corresponding to the loss of the 100 bp long exon 2. To confirm the predicted loss of ITGAL protein, flow cytometry of splenic leukocytes labelled with a fluorescently conjugated anti-ITGAL antibody was conducted. As predicted, none of the CC042 leukocyte populations examined, including CD4^+^ and CD8^+^ T cells, expressed the ITGAL protein (**Fig. 7c**). In contrast, all C57BL/6J leukocytes examined expressed the ITGAL protein.

To confirm *in vivo* the role of the *Itgal* deletion in the extreme susceptibility of CC042 strain to *S*. Typhimurium, we performed a quantitative complementation test (see raw data in Supplementary Table S3). Compound heterozygous mice carrying a KO *Itgal* allele and a CC042 deleted *Itgal* allele had significantly higher liver bacterial loads (~6.3 Log CFUs/g liver, Fig. 8a) compared with mice heterozygous for either of these two alleles (~5.1-5.3 Log CFUs/g liver). Therefore, there was no significant difference between *Itgal^+^/ Itgal^+^* and *Itgal^+^/ Itgal^KO^* mice in the B6 background (red line), while the difference between *Itgal^+^/ Itgal^del^* and *Itgal^KO^/ Itgal^del^* mice was highly significant (blue line, p = 2.1 10^-6^). These results demonstrate that the *Itgal* deletion present in CC042 mice contributes to their increased susceptibility. Remarkably, the *Itgal* KO mutation did not impact liver bacterial load in a B6 inbred background (Fig. 8b).

**Figure 8.**
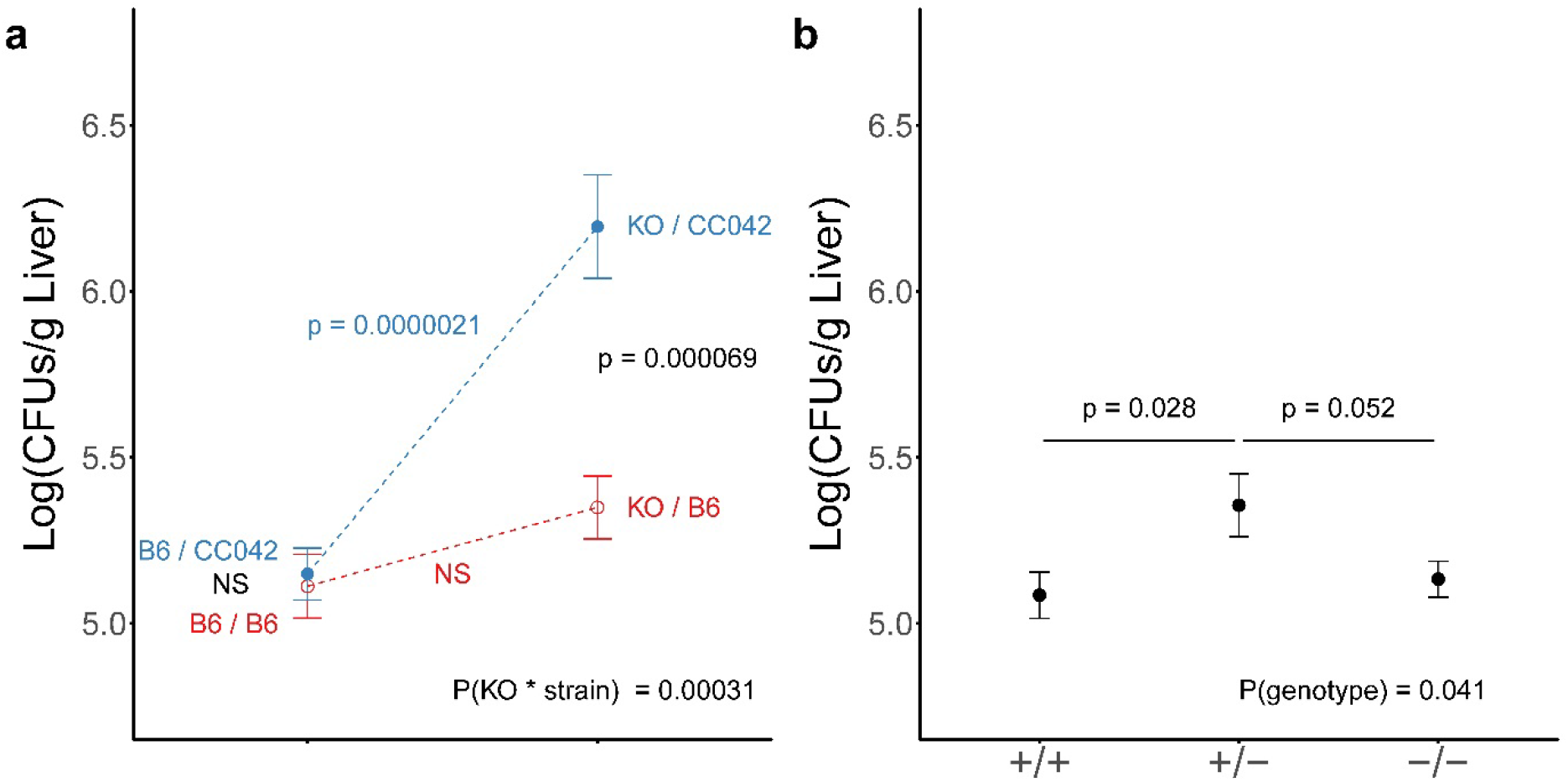
Quantitative complementation test confirmed the role of the CC042 *Itgal* loss-of-function variant in the susceptibility to *S*. Typhimurium. Bacterial load in liver (**a**) at day 4 post-infection with *S*. Typhimurium in C57BL/6J (B6/B6) (n=23), (*Itgal*^-/-^ x C57BL/6J)F1 (KO/B6) (n=17), (C57BL/6J x CC042)F1 (B6/CC042) (n=31) and (*Itgal*^-/-^ x CC042)F1 (KO/CC042) (n=20) mice with data pooled from two independent experiments. On the left are mice which carry one B6 allele and either a B6 or a CC042 allele. On the right are mice which carry one *Itgal* KO allele and either a B6 or a CC042 allele. (*Itgal*^-/-^ x CC042) F1 (KO/CC042) mice show bacterial loads higher by approximately 1 Log(CFUs) than the three other groups, indicating absence of complementation between *Itgal* KO and CC042 alleles. Differences between groups were assessed by one-way ANOVA. Interaction between genetic background (C57BL/6J or CC0042) and *Itgal* genotype (*+/+* or *-/-*) was assessed by two-way ANOVA. In an inbred C57BL/6J background (**b**), the *Itgal* KO mutation did not significantly impact the susceptibility to *S*. Typhimurium (X-axis: genotype at the *Itgal* locus; n = 34, 17 and 23, respectively; p-values from pairwise Student’s t-test).

## Discussion

The CC is a murine genetic reference population that was developed to reflect the genetic diversity and complexity of the human population [16]. In a previous study, we utilized the CC to investigate the role of genetic factors on host susceptibility to *Salmonella* Typhimurium infection [19]. The CC042 strain was identified as being highly susceptible to *S*. Typhimurium infection [19]. In this study, we sought to characterize the immunophenotype of the CC042 strain and to identify the underlying genetic factors contributing to the extreme susceptibility phenotype. We demonstrated that CC042 mice display a generalized immunodeficiency with reduced cellularity of the spleen, thymus and bone marrow. CC042 mice also exhibited alterations in immune progenitor development in both the thymus and the bone marrow. Using genetic linkage mapping and *in vivo* complementation testing we identified a *de novo* 15 bp deletion in the intron 1 of the *Itgal* gene as causatively linked to the *Salmonella* susceptibility phenotype in CC042 mice. This mutation results in exon 2 skipping and premature stop codon.

The *Itgal* mutation identified in CC042 mice was demonstrated to result in complete loss of ITGAL protein expression on all leukocytes examined. ITGAL is an integrin α-chain protein which in conjunction with the ITGB2 β-chain (ITGB2 can dimerize with any of ITGAL, ITGAM, ITGAD or ITGAX) forms the lymphocyte function associated antigen 1 (LFA-1) integrin complex [27]. The LFA-1 complex is expressed on all leukocytes and plays a critical role in various immunological functions [27]. Circulating leukocytes utilize LFA-1 to bind to its cognate ligand ICAM-1 expressed on endothelial cells [28, 29]. Binding allows leukocytes to anchor in the bloodstream prior to extravasation into lymph nodes or inflamed tissues [28, 29]. LFA-1 also enables stabilization of the immunological synapse during T cell activation, thus permitting prolonged T cell receptor (TCR):major histocompatibility complex (MHC)interactions [28, 29]. Given that LFA-1 plays a crucial role in leukocyte migration and T cell activation, it is unsurprising that absence of ITGAL function may result in ineffective immune responses to *Salmonella* infection in CC042 mice.

CC042 mice displayed significant defects in splenic, thymic and bone marrow immune cell populations. A study by Bose et al. similarly reported that *Itgal*^-/-^ mice had reduced splenic and thymic cellularity but did not observe a reduction in bone marrow cell counts [30]. Moreover, *Itgal*^-/-^ mice or mice treated with anti-LFA-1 antibodies are consistently reported to have reductions in splenic T cells [30–32]. Multiple reports specifically highlight defects in the CD8^+^ T cell subset in *Itgal*^-/-^ mice in terms of numbers, activation, proliferation and recruitment to infected tissues [31–33]. CC042 mice were similarly observed to have a more significant impairment in CD8^+^ T cells in comparison to CD4^+^ T cell populations. Defects in T cell activity may be due to the inability of T cells to sustain long-lived interactions with APCs. *Itgal*^-/-^ T cells are noted to traffic at a quicker rate through lymph nodes while T cells expressing low levels of LFA-1 have been shown to only transiently form contacts with APCs resulting in diminished expression of effector proteins including IFN-γ [34, 35]. This is consistent with our observation of low expression of IFN-γ by CD4^+^ and CD8^+^ T cells after TCR stimulation in CC042 mice. Notably, both CC042 and *Itgal*^-/-^ mice have previously been shown to display suppression of IFN-γ in lung tissue upon *M. tuberculosis* infection [36, 37]. Production of IFN-γ by T cells is crucial for the clearance of intracellular bacterial infections such as *Salmonella* as IFN-γ activates bacterial killing within infected macrophages [38, 39]. Notably, IFN-γ production has been shown to be inducible by ISG15 through an ITGAL dependent signalling mechanism [40]. Thus, it is most likely that the immunosuppressed phenotype observed in CC042 mice in response to *Salmonella* may be due to reduced T cell activation and suppression of IFN-γ production via the ITGAL dependent pathway.

Alterations in thymocyte maturation were also observed in CC042 mice potentially explaining the corresponding reduction in peripheral T cell counts. ITGAL has been implicated in thymopoiesis as *Itgal*^-/-^ mice exhibit a general reduction in all thymocytes while use of anti-ITGAL antibodies has been shown to result in impairment of double positive CD4^+^/CD8^+^ and single positive CD8^+^ T cell development [30, 41]. Thymocyte defects may be due to alterations further upstream as bone marrow hematopoiesis was also defective in CC042 mice. Hematopoietic progenitor populations have also been observed to be altered in *Itgal*^-/-^ mice [30]. In competitive reconstitution assays in irradiated mice, *Itgal*^-/-^ derived bone marrow progenitors were unable to compete against WT cells presenting a role for ITGAL in hematopoietic cell generation [30]. It should be noted that the mutation in *Kitl* also inherited from WSB/EiJ may be contributing to the defects in hematopoietic progenitor generation.

Several animal studies have delineated a significant role for ITGAL in the response to infection. *Itgal^-/-^* mice have been reported to be highly susceptible to bacterial infection by both intracellular *M. tuberculosis* and extracellular *Streptococcus pneumoniae* as characterized by reduced survival, increased bacterial burdens and defects in leukocyte recruitment to infected tissues [36, 42]. During *M. tuberculosis* infection, the absence of the ITGAL protein was shown to result in impaired containment of the bacteria demonstrated by diffuse lung granuloma formation [36]. Interestingly, upon *M. tuberculosis* infection, CC042 mice displayed significantly increased bacterial burdens of the lung and spleen, increased necrotic lung granuloma formation and more rapid disease progression compared to C57BL/6J mice [37]. This phenotype is likely partly explained by the *Itgal* deletion we have reported here. Studies utilizing anti-LFA-1 antibodies to neutralize ITGAL protein activity have shown that ITGAL function is necessary for timely clearance of *respiratory syncytial virus* (RSV) and for control of parasitemia during *Trypanosoma cruzi* infection [33, 43]. In contrast, two separate studies have reported that *Itgal*^-/-^ mice display increased resistance to *Listeria monocytogenes* infection [31, 44]. Counterintuitively, resistance to *L. monocytogenes* occurs despite *Itgal*^-/-^ mice displaying a significant reduction in CD8^+^ T cell numbers and lytic activity during *L. monocytogenes* infection [31]. The discrepancy in response to *L. monocytogenes* as compared to other pathogens may be due to differential methods of leukocyte recruitment in response to various inflammatory stimuli. One study found that neutrophil recruitment during *S*. Typhimurium and *L. monocytogenes* infection operates in ITGB2 dependent and independent manners respectively [45].

To our surprise, mice homozygous for the *Itgal* KO mutation on the B6 inbred background did not show increased liver bacterial load. However, increased susceptibility was observed in compound heterozygous mice on a B6/CC042 F1 background. These results suggest that the impact of ITGAL loss of function may depend on the genotype at other loci. There are in fact many examples were the phenotype induced by the inactivation of a gene is influenced by the genetic background [46, 47]. One of the modifier loci could be *Stsl6* which we identified in the F2 cross. While the comparison of the founder haplotypes in CC042 and other CC strains suggested that, like for *Stsl7*, the causative variant was likely a *de novo* mutation proper to CC042, we failed to identify a candidate private variant for *Stsl6* from the published data. Moreover, due to the linkage between these two loci, the F2 did not allow to investigate genetic interactions between the two loci.

One of the main advantages of the CC is the ability to model complex traits in genetically diverse populations [17], in particular for studying host-pathogen interactions [18]. The hope is that genetic factors uncovered through CC mouse studies may be applicable to studies of human disease [48]. Indications that ITGAL may be important for responses to *Salmonella* infection in humans is supported by the finding that ITGAL is upregulated upon infection with *S*. Typhi in human volunteers [49]. Moreover, ITGAL has been implicated in a range of inflammatory diseases in humans. A recent genome wide association study identified an ulcerative colitis risk allele at the *Itgal* locus resulting in upregulation of ITGAL protein expression [50]. ITGAL has also shown to be upregulated in systemic lupus erythematosus likely due to reported DNA hypomethylation at the ITGAL promoter [51, 52]. Meanwhile, gene pathway analysis has uncovered a multiple sclerosis susceptibility gene network involving both *Itgal* and its cognate receptor gene *Icam1* [53]. Furthermore, deficiency of ITGAL’s dimerization partner, ITGB2, results in leukocyte adhesion deficiency I (LAD-I) which is classically characterized by recurrent bacterial infections [54]. These studies suggest that overexpression of ITGAL may result in excessive activation of the immune system resulting in inflammatory disease while deficiency may result in immunosuppression.

Characterization of the CC042 mouse strain has provided a foundation for future use of the CC in infection susceptibility studies. We have used this model to identify and characterize the novel *Salmonella* susceptible CC042 strain and to identify *Itgal* as a gene critical in the *Salmonella* response pathway.

## Material and methods

### Ethics statement and animals

Animal experiments performed at McGill University were conducted in accordance to guidelines provided by the Canadian Council on Animal Care (CCAC). Guidelines include the Guide to Care and use of experimental animals, vol 1, 2nd edition, choosing an appropriate endpoint in experiments using animals for research, teaching and testing, euthanasia of animals used in science, husbandry of animals in science, laboratory animal facilities – characteristics, design and development, and training of personnel working with animals in science. The animal-use protocol was approved by the McGill University Facility Animal Care Committee (protocol no. 5797). Animal experiments performed at the Institut Pasteur were conducted in compliance with French and European regulations and were approved by the Institut Pasteur Ethics Committee (project #2014-0050) and authorized by the French Ministry of Research (decision #8563).

CC042/GeniUnc (CC042) mice were originally generated through the CC at Geniad [20] in Australia and then maintained at the Institut Pasteur (Paris) and McGill University (Montréal) under specific-pathogen-free conditions. C57BL/6J (B6) mice were purchased from the Jackson Laboratory (stock #000664). C57BL/6NCrl (B6N) mice used to generate (B6NxCC042)F2s were purchased from Charles River. B6.129S7-*Itgal^tm1Bll^*/J (*Itgal* KO) mice used to perform complementation tests were purchased from the Jackson Laboratory (stock #005257). F1s were obtained by crossing CC042 males with B6N females and were intercrossed to produce F2s.

### Salmonella Typhimurium infection

The infectious dose was generated via culture of frozen *S*. Typhimurium strain SL1344 stocks in trypticase soy broth at 37°C until an optical density of 0.1-0.2 at 600 nm was reached. To determine the specific bacterial concentration, the bacterial suspension was diluted in saline and plated on trypticase soy agar (TSA). Using the resulting concentration, an inoculum of 1000 CFU in 200 μL was administered to mice via intravenous injection into the caudal vein. All healthy mice aged between 7-12 weeks were included. Investigators were blinded to genotypes during the monitoring of *Salmonella*-infected mice. The dose was verified via serial dilution of the inoculum and bacterial culture on TSA. Following infection, mice were monitored and assessed using body condition scoring. Mice were humanely euthanized at different time points post *Salmonella* infection for sample collection. To determine bacterial loads, the spleen and liver were aseptically removed, weighed and added to 0.9% saline. The organs were homogenized using a Polytron homogenizer (Kinematica, Bohemia, NY), serially diluted in PBS and cultured on TSA.

### Hematology

For complete blood count and white blood cell differential, blood samples were collected in EDTA tubes from mice aged 8-10 weeks. Analyses were performed at the Comparative Medicine Animal Resources Centre, McGill University.

### Histology

Tissues were collected and placed in 10% formalin at room temperature for 24 hours to enable fixation. The organs were then transferred to 70% ethanol solution at 4°C followed by organ processing, embedding and sectioning at the Goodman Cancer Research Center histology facility, McGill University. The resulting tissue sections were stained with hematoxylin and eosin prior to microscopic examination.

### Flow cytometry analyses

For flow cytometry analyses, whole spleens and thymi were aseptically collected, weighed and transferred to 3 mL of phosphate buffered saline (PBS). Tissues were mechanically dissociated using the backend of 3 ml syringes into a 70 μm strainer (Fisherbrand). The resulting cell suspensions were passed through a second 70 μm strainer and transferred to a 15 mL conical tube. The cells were spun at 1400 revolutions per minute (RPM) for 5 minutes at 4°C. The supernatant was discarded and the pellets were resuspended in 10 mL (spleen) or 5 mL (thymus) of ammonium-chloride-potassium (ACK) lysis buffer at room temperature. The suspension was spun at 1400 RPM for 5 minutes at 4°C and the supernatant was discarded. The cell pellets were resuspended in 10 mL PBS and passed through a third cell strainer to generate single cell suspensions. For bone marrow cells preparations, femurs were aseptically collected. The bones were cleaned of flesh using a scalpel blade. The epiphyses of the femurs were cut and a 25 G needle and syringe was used to pass 2 mL of PBS through the bone to displace the bone marrow. The bone marrow containing media was transferred to a 50 mL tube and 2 mL of red blood cell lysis buffer (Sigma Aldrich) was added. After 1 minute of gentle mixing, 15 mL of PBS was added and the tubes were centrifuged for 7 minutes at 1400 RPM and the supernatant discarded. The resulting cell pellet was resuspended in 2 mL of PBS.

The following monoclonal antibodies were used for flow cytometric analysis of spleen, thymus and bone marrow samples. Invitrogen: CD4-PE-Cy7 (GK1.5), CD11b-APC (M1/70), Ly6C-PE (HK1.4). BioLegend: B220/CD45R-PerCP-Cy5.5 (RA3-GB2), CD8α-APC-Cy7 (53-6.7), CD11a-AlexaFluor488 (M17/4), CD49b-pacific blue (DX5), c-Kit-PE-Cy7 (2B8), F4/80-PE-Cy7 (BM8), Flt3-PE (A2F10), IL7R-PE (A7R34), Ly6G-APC-Cy7 (1A8), Sca-1-APC-Cy7 (D7), TER119-PerCP-Cy5.5 (TER-119). eBioscience: B220/CD45R-APC (RA3-6B2), B220/CD45R-PerCP-Cy5.5 (RA3-GB2), CD11b-PerCP-Cy5.5 (M1/70), CD11c-eFluor450 (N418), CD16/CD32-FITC (93), CD25-AlexaFluor488 (eBio3C7), CD34-eFluor450 (RAM34), CD44-PE (IM7), CD48-HM48.1 (FITC), CD69-FITC (H1-2F3), CD150-alexafluor647 (mShad150), MHCII-FITC (AF6-120.1), Sca-1-APC (D7), TCRβ-PE-Cy5 (H57.597).

Live cells in cell suspensions prepared from spleen, thymus and bone marrow were counted using a hematocytometer, plated in 96 well plates and washed twice with PBS. The cells were stained with 1:400 Zombi Aqua Fixable viability dye (BioLegend) for 15 minutes at room temperature in the dark followed by washing with PBS. Cells were incubated with antibody for 20 minutes at 4°C. The cells were fixed using Cytofix/Cytoperm (BD) according to the manufacturer’s directions, washed in PBS and resuspended in PBS containing 2% fetal bovine serum (FBS). Cell acquisition was completed using the FACS canto II (BD) or LSR Fortessa cell analyzer (BD) flow cytometers (McGill Flow Cytometry Core Facility) using FACs Diva software (BD). Compensation was completed using OneComp eBeads (Invitrogen) and data was analyzed using FlowJo software version 10.0.8r1.

For cytokine intracellular staining, splenocytes were stimulated with pre-coated anti-CD3 (5μg/ml, eBioscience) and soluble anti-CD28 (2μg/ml, eBioscience) in the presence of Protein Transport Inhibitor Cocktail (eBioscience) for 6 hours at 37°C and 5% CO2 in complete RPMI (10% FBS, 1mM sodium pyruvate, 1 Non-essential amino acids, 1% P/S, and β-mercaptoethanol). Cells were incubated with the following antibodies: α-CD4 PE (GK1.5), α-CD8α BV421 (53-6.7, BioLegend), and α-TCRβ FITC (H57-597). Cells were then fixed and permeabilized as per manufacturer’s protocol (Cytofix/Cytoperm, BD) and stained intracellularly with the following antibodies: α-TNFα PerCP-Cy5.5 (MPG-XT22, BioLegend) and α-IFNγ APC (XMG1.2) for 30 min at 4°C. Cells were acquired on an eight-color FACSCanto II using FACS Diva software (BD). The data was analyzed using FlowJo version 10.0.8r1software. Doublets were removed by SSC-H versus SSC-W gating.

### Bone marrow derived macrophage (BMDM) cell culture, RNA extraction, reverse transcription, PCR and gel electrophoresis

Bone marrow was extracted from mouse femurs as described above (flow cytometry analyses). The 2 mL cell suspension that was recovered was split into two petri dishes and 7 mL of complete RPMI (RPMI, 10% FBS, 1 % pen/strep) and 4 mL of macrophage colony stimulating factor (M-CSF) was added to each dish. The cells were incubated at 37°C at 5% CO2. At day 3 of culture, the supernatant was removed and replaced with 6 mL RPMI. The dishes were scraped and 10 mL of RPMI and 8 mL of M-CSF was added. The suspension was split into 2 petri dishes. At day 6, the supernatant was removed and the cells were rinsed once in 5 mL PBS before being collected in 5 mL of PBS. The cells were centrifuged for 7 minutes at 1400 RPM. The supernatant was discarded and the pellets were resuspended in TRIzol (Invitrogen, Cat #15596-018) for RNA extraction. Reverse transcription of 2.5 μg of RNA was conducted using the ABM 5X All-In-One RT MasterMix (Cat #G486) and amplification of the resulting cDNA was completed using a standard PCR reaction using EasyTaq DNA polymerase (Transgen Biotech, Cat #AP111-01) (forward primer: 5’ CCATGCAAGAGAAGCCACCAT 3’, reverse primer: 5’ AGTGATAGAGGCCTCCCGTGT 3’). RNA was visualized using gel electrophoresis on 2% agarose gels.

### F2 cross and QTL mapping

One hundred and ninety-six offspring from an F2 cross of CC042 and C57BL/6NCrl mice were infected with *Salmonella* Typhimurium and euthanized at day 4 post-infection. Spleen and liver bacterial loads were determined as described above. Ninety-four animals with extreme phenotypes (half with high and half with low bacterial loads) were selected for genotyping. SNP genotyping was performed by Neogen Inc. (Lincoln, NE), using the Mouse Universal Genotyping Array (MUGA) containing 7.8K SNPs [21]. All statistical tests (Pearson’s R correlation coefficient, QTL mapping) were performed using R statistical software. QTL mapping on liver bacterial load was performed by using the J/qtl interface 1.3.5 for R/qtl software version running under R 3.2.2 [22]. Significance thresholds of LOD scores were estimated by 10,000 permutations of experimental data.

### CC042 private variants in the QTL regions

Most CC strains were recently sequenced [23] and data are publicly available, including the list of private variants specific to each CC strain. We retrieved the CC042 private variants localized within the *Stsl6* and *Stsl7* QTLs. Three CC042 private variants were present within *Stsl6* (*Ush1c, Ccdc123, 4930435C17Rik*) and only one within *Stsl7* (*Itgal*).

### Itgal genotyping

Amplification of the region containing the CC042 *Itgal* deletion was conducted using a standard polymerase chain reaction (PCR) (forward primer: 5’ TGCTTGGGTGTAGGCAGCCTCA 3’, and reverse primer: 5’ CTTCAATCTGCAAGACCTGGTA 3’). DNA amplicons were digested using FauI with CutSmart buffer (New England BioLabs, R0651S) for 4 hours at 55°C. The reaction was stopped by incubating the samples at 80°C for 15 minutes. The digested DNA was run on 1.5% agarose gel in tris-borate-EDTA (TBE) buffer.

### Quantitative complementation testing

CC042 mice were crossed with C57BL/6J and *Itgal* KO mice to produce *Itgal^B6^/Itgal^CC042^* and *Itgal^KO^/Itgal^CC042^* genotypes on a B6/CC042 genetic background. C57BL/6J were crossed with *Itgal* KO mice to produce *Itgal^B6^/Itgal^KO^* genotype on a B6 genetic background. Mice were infected with *S*. Typhimurium and bacterial loads were determined at day 4 post-infection as described above.

## Data availability

All relevant data that support the findings of this study are within the paper and its supporting information files.

## Acknowledgements

We thank Line Larivière and Hyejin Park for technical assistance and Patricia D’Arcy (Mc Gill), Isabelle Lanctin and Tommy Penel (DTPS/C2RA-Central Animal Facility, Institut Pasteur) for mouse breeding and screening. D.M. is a recipient of a Canadian Institutes of Health Research (CIHR) project grant (MOP133700) and a Natural Sciences and Engineering Research Council (NSERC) discovery grant (RGPIN-2017-04717). A.N. is a Canada Research Chair Tier II in Hematopoiesis and Lymphocyte Differentiation and a recipient of CIHR project grant PJT-153016 and NSERC discovery grant RGPIN-2016-05657”. Funding had been provided to M.T. by the NSERC - Undergraduate Student Research Award and the Fonds de Recherche du Québec Nature et Technologies and to J.K. by the McGill Faculty of Medicine - Solvay Fellowship. The flow cytometry work/ cell sorting was performed in the Flow Cytometry Core Facility for flow cytometry and single cell analysis of the Life Science Complex (McGill University) and supported by funding from the Canadian Foundation for Innovation. This study was also supported by AgroParisTech (France) through a PhD fellowship to J.Z.

---

**Figure S1.**
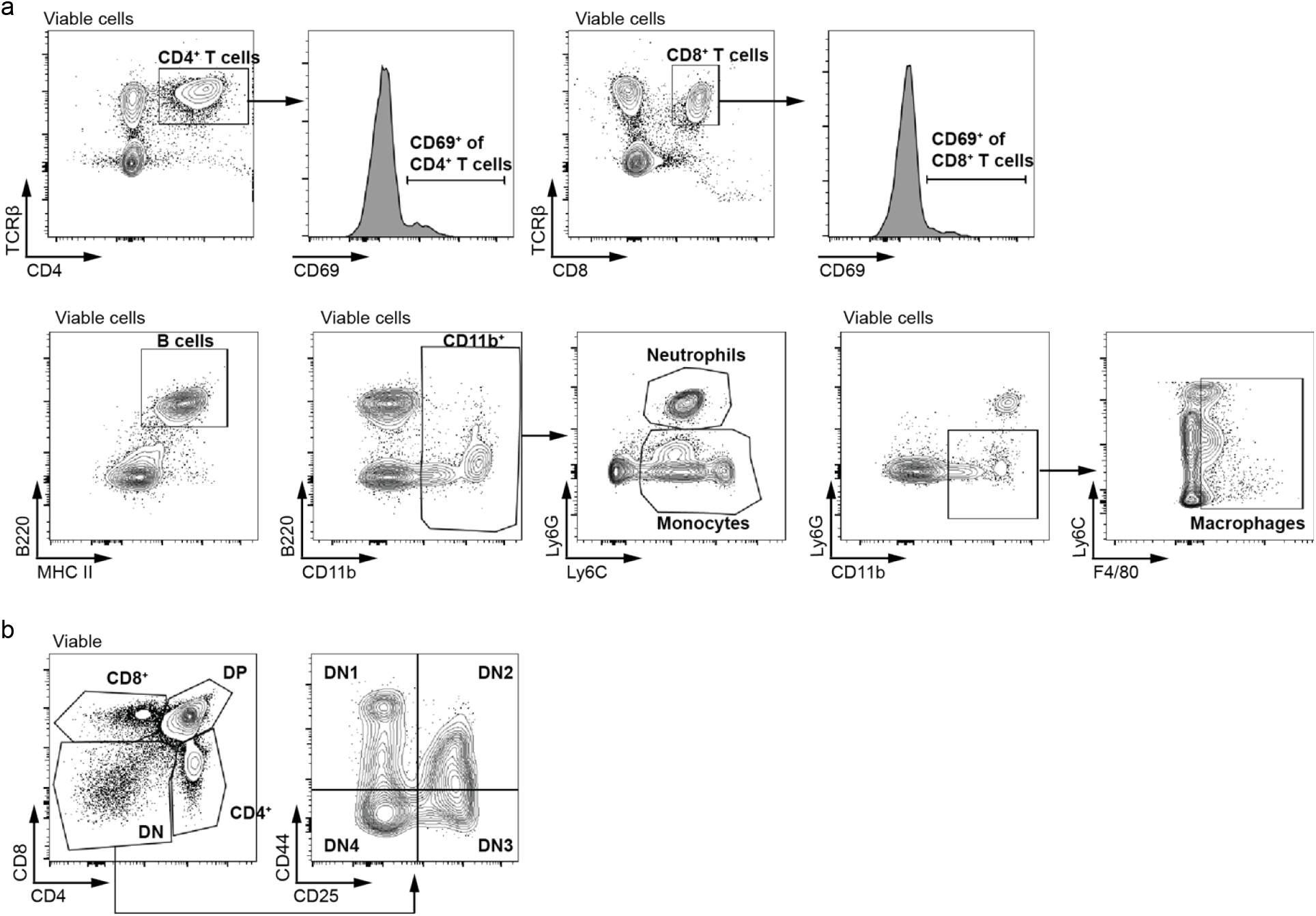
Representative flow cytometry gating schemes used for analysis of spleen and thymus. Plots show gating used for spleen (**a**) and thymus (**b**) samples where cells shown for each plot are derived from within the gated region of the previous plot.

**Figure S2.**
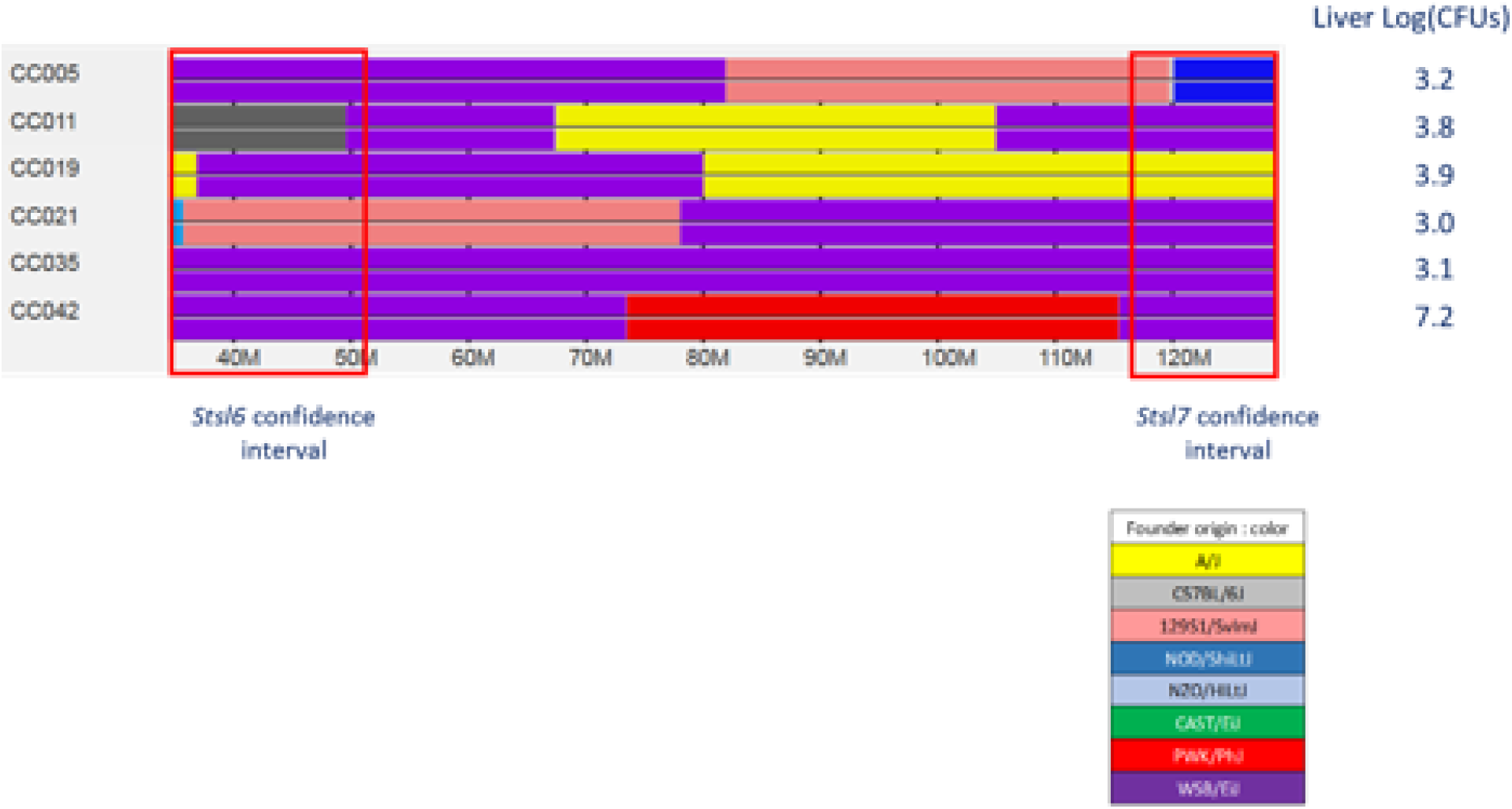
Susceptibility to *S*. Typhimurium of CC strains with a WSB/EiJ haplotype at either Chr 7 QTLs. Schematic representation of the haplotype structure of CC strains carrying WSB alleles at either *Stsl6* or *Stsl7 or both* QTLs. Corresponding bacterial load in liver (LOG10) at day 4 post-infection with *S*. Typhimurium (data from Zhang et al. 2018) is shown on the right. The 1.5 LOD confidence intervals of the QTLs are boxed in red. CC035 mice carry the same WSB/EiJ-derived haplotype as CC042 but show much lower bacterial load in the liver.

**Table S1 | Raw data for Parental, (C57BL/6N x CC042)F1 and (C57BL/6N x CC042)F2 mice.** For each animal, Strain, Sex, Age, Spleen and Liver bacterial loads are provided.

**Table S2 | Raw data for the subset of 94 (C57BL/6N x CC042)F2 mice with extreme bacterial loads used for QTL mapping.** For each animal, Id, Experiment number, Sex, Age, Spleen and Liver bacterial loads are provided. Genotyping results (MUGA array) are also provided.

**Table S3 | Raw data for animals used in the *Itgal* quantitative complementation test.**

